# Mutation of *OsCDS5* confers broad-spectrum disease resistance in rice

**DOI:** 10.1101/2023.12.18.572258

**Authors:** Qiping Sun, Yongxin Xiao, Le Song, Lei Yang, Yin Wang, Wei Yang, Qun Yang, Kabin Xie, Meng Yuan, Guotian Li

## Abstract

Phospholipids are important components of biological membranes, participating in various biological processes, including plant development and responses to biotic and abiotic stresses. A previous study showed that mutation of the rice *OsCDS5* (<u>C</u>DP-<u>D</u>AG <u>S</u>ynthase) gene alters lipid metabolism, causing enhanced abiotic stress responses, yellowing of leaves at the seedling stage and delayed plant development. Here, we observed that the *Oscds5* mutant shows enhanced resistance to rice blast, bacterial blight and bacterial leaf streak. Mutation of *OsCDS5* promotes production of reactive oxygen species (ROS) and increases the expression level of multiple defense-related genes. Transcriptomic analyses indicate that genes involved in responses to stress, biotic/abiotic stimuli and metabolic processes are highly upregulated and enriched in mutant *Oscds5*. Metabolomic analyses show that differential metabolites are enriched in the lipid metabolic and tryptophan metabolic pathways. The decreased level of phosphatidylinositol (PI) and increased level of serotonin likely contribute to enhanced disease resistance of the *Oscds5* mutant. Taken together, mutation of *OsCDS5* enhances abiotic and biotic stress responses, and *OsCDS*5 may be a promising target in genetic engineering to enhance the resilience of rice to abiotic and biotic stresses simultaneously.

---

As the world’s population grows, more food is needed. However, biotic and abiotic stresses, such as devastating crop diseases, drought and salinity, pose serious threats to global food security (Dhankher & Foyer, 2018). Among crop diseases, rice blast alone, which is caused by the filamentous fungus *Magnaporthe oryzae*, results in global losses of rice grains equivalent to food for 60 million people annually (Nalley et al., 2016). In addition to rice blast, bacterial diseases such as bacterial blight and bacterial leaf streak of rice caused by *Xanthomonas oryzae* pv. *oryzae* (*Xoo*) and *Xanthomonas oryzae* pv. *oryzicola* (*Xoc*), respectively, occasionally cause severe grain losses up to 32%-50% of the total yield (Li et al., 2019, Liu et al., 2014, Liu et al., 2021). Thus, crop cultivars resilient to biotic and abiotic stresses are key to fulfilling the increasing demand for the world supply. Identification of common targets for biotic and abiotic stress responses lays the foundation for developing crops resilient to biotic and abiotic stresses simultaneously.

Phospholipids are important molecules in plants, being the components of biological membranes and signaling molecules (Jennings & Epand, 2020). Cytidine diphosphate diacylglycerol (CDP-DAG) synthase (CDS) is one of the key enzymes in phospholipid biosynthesis. CDS catalyzes the biosynthesis of CDP-DAG from phosphatidic acid (PA). PA is an important signaling molecule and the precursor of all glycerophospholipids. CDP-DAG is used to biosynthesize multiple other important phospholipid compounds, including cardiolipin (CL), phosphatidylglycerol (PG), PI, and phosphatidylserine (PS) (Yang et al., 2018). PI is the precursor of phosphatidylinositol (3) phosphate (PI3P) and phosphatidylinositol (4) phosphate (PI4P), and PI4P can be converted into phosphatidylinositol (4,5) bisphosphate (PI(4,5)P2) by adding an additional phosphate group (Blunsom & Cockcroft, 2020). A previous study showed that PI(4,5)P2, as a disease-susceptibility factor, plays important roles in powdery mildew infection and is recruited to the extrahaustorial membrane during fungal-*Arabidopsis thaliana* interactions (Qin et al., 2020).

In plants, the CDP-DAG synthase gene family has expanded. In the budding yeast *Saccharomyces cerevisiae*, there is only one *CDS1* gene (Shen et al., 1996). In the model plant *A. thaliana*, there are five *CDS* genes, named *AtCDS1* to *AtCDS5* (Haselier et al., 2010). AtCDS1, AtCDS2 and AtCDS3 are localized in the endoplasmic reticulum (ER), while AtCDS4 and AtCDS5 are localized in plastids (Zhou et al., 2013). The *Atcds1* and *Atcds2* single mutants express no significant changes in growth and lipid metabolism, indicating that AtCDS1 and AtCDS2 function redundantly. In contrast, the *Atcds1 Atcds2* double mutant dies two weeks post-germination. The *Atcds1 Atcds2* double mutant contains slightly smaller chloroplasts, and accumulates more PA, 8-fold higher than that of the wild-type (WT) plants (Zhou et al., 2013). Similarly, the *Atcds4 Atcds5* double mutant is defective in thylakoid development and has a 40% reduction in PG content compared to the WT (Haselier et al., 2010). Rice contains at least three CDP-DAG synthases homologous to yeast Cds1. RBL1, the CDP-DAG synthase with the highest expression level among the three CDS genes in rice leaves, regulates cell death and plant immunity (Sha et al., 2023). The *rbl1* mutant shows broad-spectrum disease resistance to rice blast, rice false smut and bacterial blight, and shows higher levels of reactive oxygen species (ROS), salicylic acid (SA) and defense-related genes than the WT. The levels of PG, PI and PI(4,5)P2 are decreased in the *rbl1* mutant. Overexpression of rice phosphatidylinositol synthase gene *PIS1* (*OsPIS1*) increases the level of PI and disease susceptibility in *rbl1*. PI(4,5)P2, also as a susceptible factor in rice, is enriched near the infection site of *M. oryzae*. Decreased levels of PI and PI(4,5)P2 are not conducive to *M. oryzae* infection, endowing rice with disease resistance. Recently, an additional predicted *CDS* was identified in rice, named *OsCDS5*. OsCDS5 is localized to the chloroplasts and ER (Hong et al., 2018). In the *Oscds5* mutant, the thylakoid membrane is defective, and the leaves are pale at the seedling stage, resulting in delayed growth. Mutation of *OsCDS5* also results in PA accumulation and decreased levels of PG and PI. Importantly, mutation of *OsCDS5* confers hyperosmotic tolerance in rice. However, the role of *OsCDS5* in biotic stress responses is unknown.

To investigate whether *OsCDS5* is involved in disease resistance in rice, we inoculated the mutant *Oscds5* with pathogens including *M. oryzae, Xoo* and *Xoc*. Rice plants used for pathogen inoculation, which are in the Dongjin (*Japonica*) background, were grown at the experimental station under natural long-day conditions in the summer. For *M. oryzae* infection, leaves of 6-week-old rice plants were wounded with a mouse ear punch, and a mycelial block (2 mm x 2 mm) from fresh *M. oryzae* cultures was applied to the wounded area, which was then sealed with cellophane tapes (Li et al., 2017). Three *M. oryzae* strains LN6, LN10 and LN13 were used in the infection assays. The disease symptoms were examined at 12 days post-inoculation (dpi) and the lesion areas were analyzed with software imageJ. We observed that the lesion area in mutant *Oscds5* plants was smaller than that in the WT for all the three *M. oryzae* strains (Figure 1a). More precisely, the blast lesion areas in mutant *Oscds5* were 28%, 37% and 21% smaller compared to the WT for strains LN6, LN10 and LN13, respectively. We used DNA-based quantitative PCR to analyze relative fungal biomass in the lesion, which was represented by the ratio of the *MoPot2* gene of *M. oryzae* to the *Ubiquitin* gene of rice (Kawano et al., 2010). The results showed that fungal biomass in mutant *Oscds5* was 70% and 50% lower than that in the WT for *M. oryzae* strains LN6 and LN10, respectively (Figure 1b). Taken together, these results demonstrate that mutant *Oscds5* shows enhanced resistance to three rice blast fungal strains.

**FIGURE 1.**
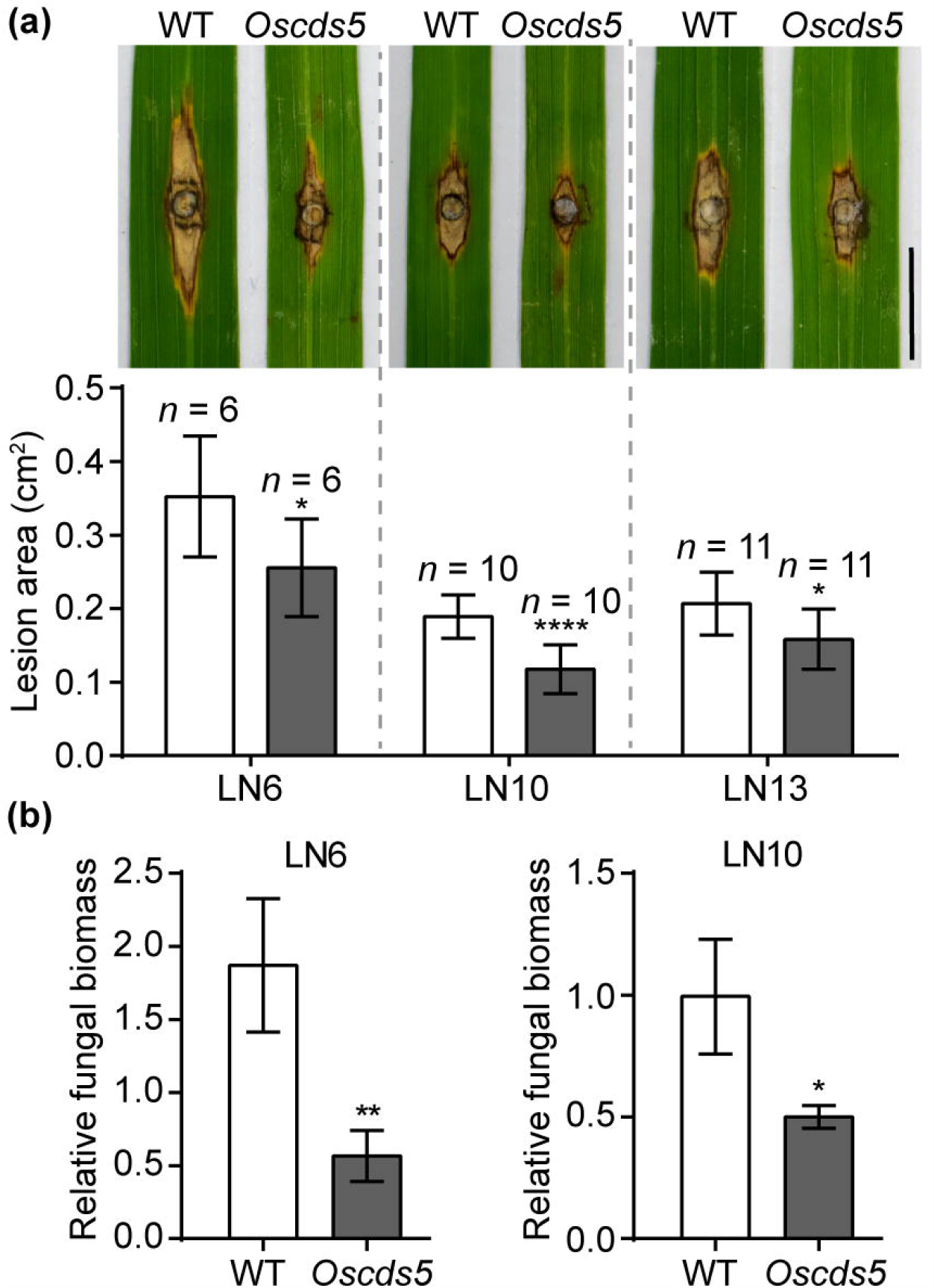
Mutation of *OsCDS5* confers resistance to rice blast. (a) Disease symptoms of the wild-type (WT) and *Oscds5* plants infected with *M. oryzae* strains LN6, LN10 and LN13 at 12 days post-inoculation (dpi). Leaves of 6-week-old rice plants were used in the punch-inoculation assays. Bar, 1Ccm. (b) Relative fungal biomass of the WT and*Oscds5* lines infected with *M. oryzae* strains LN6 and LN10 as in (a) (*n* = 3). *n* indicates the number of biological replicates used in the analyses. Asterisks indicate significant differences using the unpaired Student’s *t*-test (\**p* < 0.05, \*\**p* < 0.01, \*\*\*\**p* < 0.0001).

To investigate the resistance of mutant *Oscds5* to bacterial diseases of rice, we inoculated the *Oscds5* mutant plants at the 11-week-old stage with *Xoo* strain PXO99 by the leaf-clipping method (Ke et al., 2017, Liu et al., 2023). The disease symptoms were examined at 14 dpi (Figure 2a). The average lesion length in the *Oscds5* mutant was 3.6 cm, compared to the WT and complementation lines at 9.8 cm and 10.9 cm, respectively (Figure 2b). Similarly, in bacterial biomass assays with DNA-based quantitative PCR, the bacterial biomass in mutant *Oscds5* was 88% lower than that in the WT (Figure 2b). These results show that mutation of *OsCDS5* confers enhanced resistance to rice bacterial blight. Similarly, enhanced resistance to bacterial leaf streak caused by *Xoc* was observed for mutant *Oscds5*. In the punch inoculation assay with the *Xoc* strain HB8 (Yang et al., 2023) with the needleless syringe, mutant *Oscds5* showed disease lesions with an average length of 1.4 cm at 14 dpi, compared to the WT and complementation lines both at approximately 3.1 cm (Figure 2c). The bacterial biomass of mutant *Oscds5* was 31% lower than that in the WT (Figure 2d). These results together show that mutant *Oscds5* has enhanced resistance to bacterial blight and bacterial leaf streak.

**FIGURE 2.**
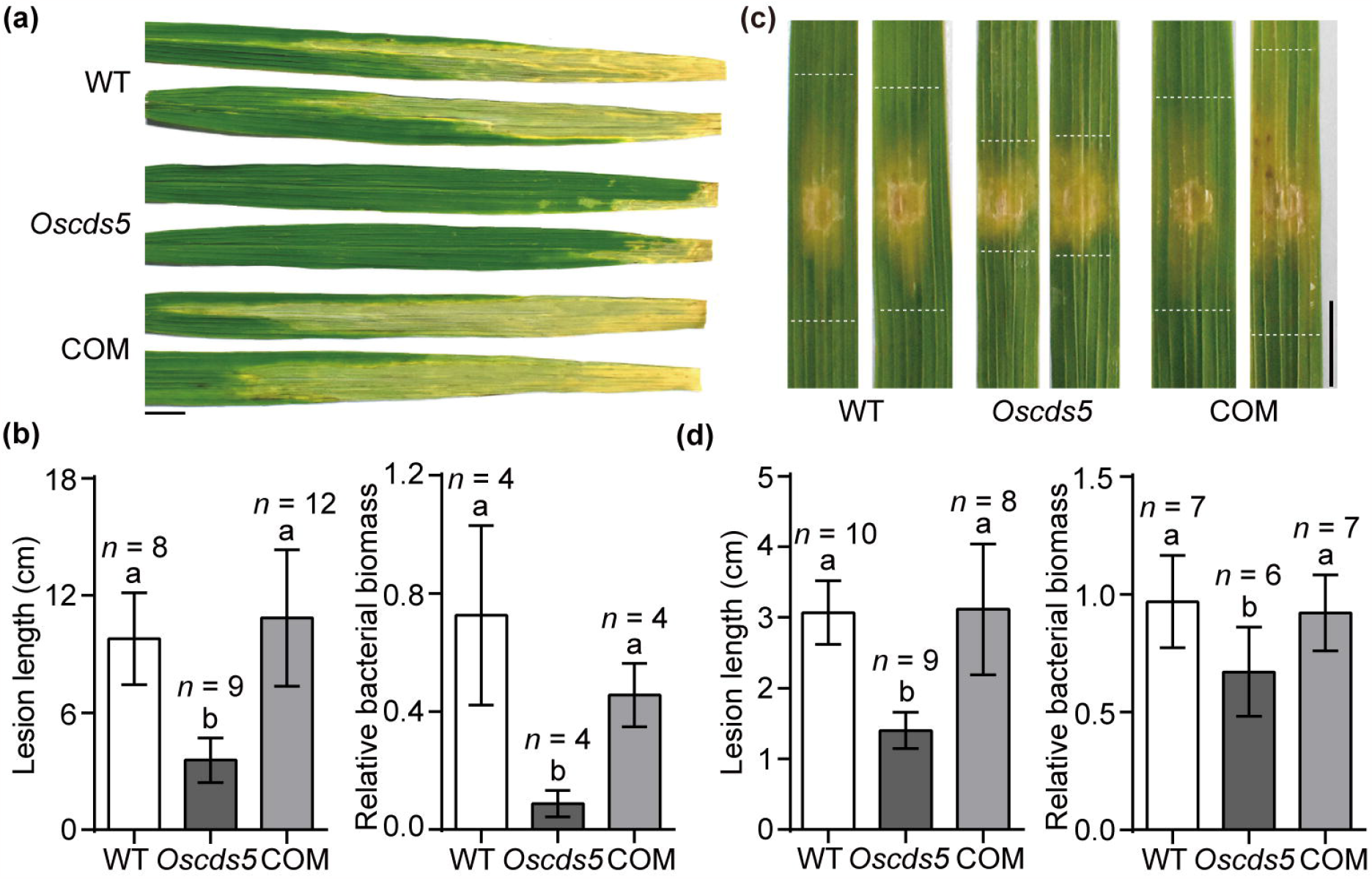
Mutation of *OsCDS5* confers resistance to rice bacterial blight and bacterial leaf streak. (a) Disease symptoms of the WT, *Oscds5* and complementation (COM) plants infected with *Xanthomonas oryzae* pv. *oryzae* (*Xoo*) strain PXO99 at 14 dpi. Flag leaves of plants at the booting stage were inoculated with the *Xoo* suspension at OD600 = 0.5 using the leaf-clipping method. Bar, 1 cm. (b) Lesion length and relative bacterial biomass of the WT, *Oscds5* and COM lines as in (a). (c) Disease symptoms of the WT, *Oscds5* and COM plants infected with *Xanthomonas oryzae* pv. *oryzicola* (*Xoc*) strain HB8 at 14 dpi. The dashed line indicates the margin of the lesion. Leaves of 5-week-old rice plants were used in infection assays. Bar, 1cm. (d) Lesion length and relative bacterial biomass of the WT, *Oscds5* and COM lines as in (c). *n* indicates the number of biological replicates used in the analyses. Different letters in the graphs denote statistically significant differences (ANOVA, *p* < 0.05).

ROS play an important role in plant immunity. To investigate the role of ROS in the enhanced disease resistance of mutant *Oscds5*, we assayed ROS production by challenging the *Oscds5* plants with chitin and flg22, pathogen-associated molecular patterns (PAMPs) typical of fungal and bacterial pathogens, respectively, which induce PAMP-triggered immunity (PTI) (Tang et al., 2021, Yang et al., 2022). In the assay, leaves of 5-week-old rice plants were cut into small disks (3 mm x 3 mm), which were gently transferred to distilled water in a 96-well plate. The sample plate was incubated under consistent light overnight, and then treated with chitin at 8 nM or flg22 at 100 nM. The chemiluminescence was immediately measured. The ROS levels in mutant *Oscds5* were 2- and 3.7-fold higher than that in the WT for chitin and flg22, respectively. And the ROS levels in the complementation line (COM) are similar to that in the WT (Figure 3a, b), suggesting that the *Oscds5* mutant shows enhanced PTI. In reverse transcription quantitative PCR (qRT-PCR) assays with 5-week-old plants, we observed 2-, 0.8-, 6.5-, and 9.5-fold higher expression levels for defense-related genes *OsAOS2, OsPAL1, OsPR1a*, and *OsWRKY70*, respectively, in the mutant compared to the WT (Figure 3c, and Table S1). And there is no significant difference in the expression levels of these disease resistance related genes between the COM and WT lines. *OsAOS2* encodes an allene oxide synthase, which participates in the jasmonic acid biosynthetic pathway. Overexpression of *OsAOS2* enhances resistance to *M. oryzae* (Mei et al., 2006). *OsPAL1* is a phenylalanine ammonia-lyase gene, important for lignin biosynthesis and plant immunity, and mutation of *OsPAL1* attenuates resistance to bacterial blight, sheath blight and rice blast (Tonnessen et al., 2015). OsPR1a and OsWRKY70 are common positive regulators of rice immunity (Choi et al., 2020, Ma et al., 2022, Song et al., 2010). These results show that basal immunity is enhanced in mutant *Oscds5*.

**FIGURE 3.**
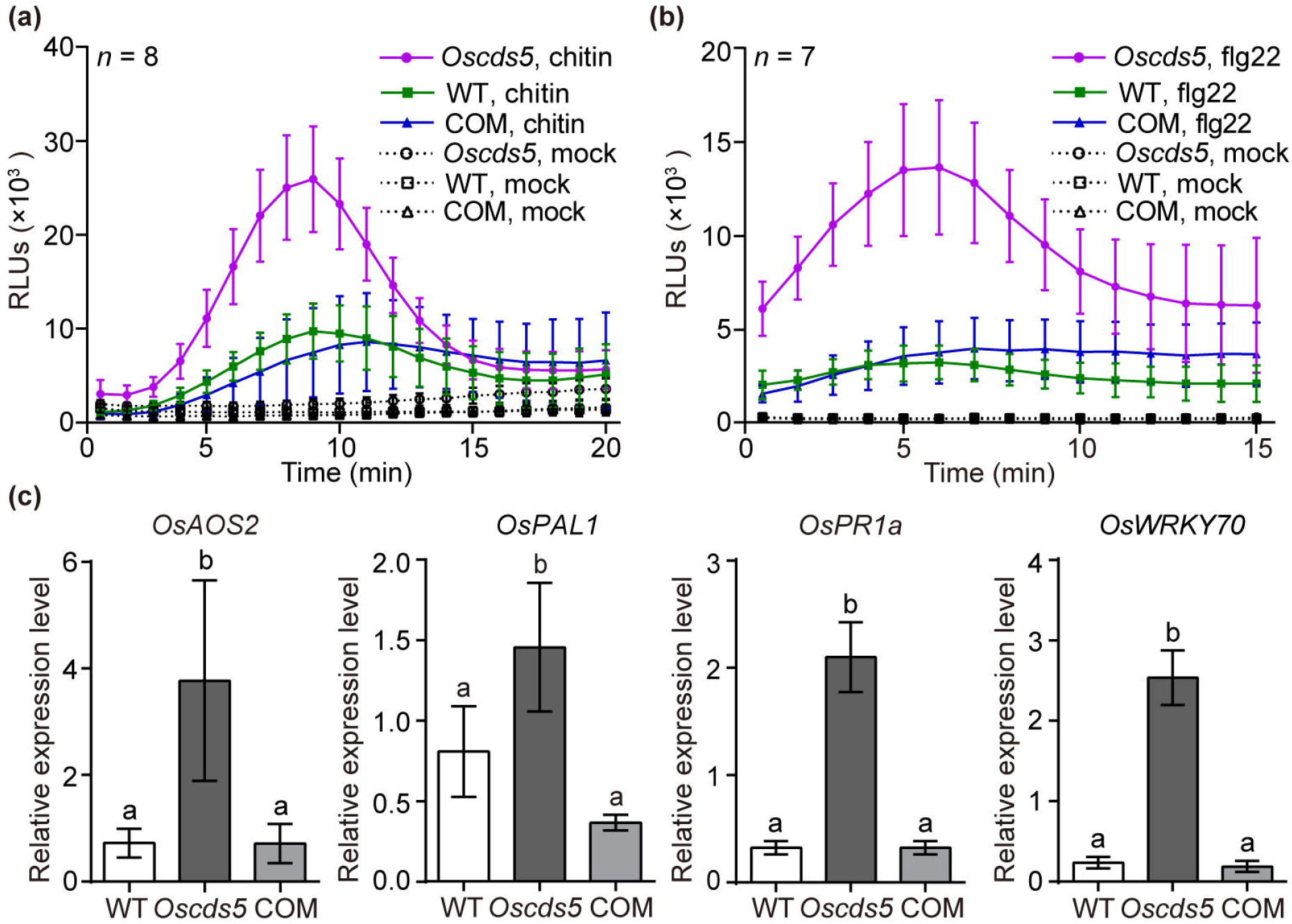
Mutation of *OsCDS5* enhances immune responses of rice. Reactive oxygen species (ROS) production in the WT, *Oscds5* and COM plants when challenged with pathogen-associated molecular patterns (PAMPs) chitin (a) and flagellin22 (flg22) (b), respectively. Leaves of 30-day-old rice plants were used in the assays. Water was used in the mock treatment. RLU, relative light unit. (c) qRT-PCR assays of defense-marker genes in the WT, *Oscds5* and COM plants (*n* ≥ 3). *n* indicates the number of biological replicates used in the analyses. Different letters on the graphs denote statistically significant differences (ANOVA, *p* < 0.05).

To further investigate the role of OsCDS5 in plant immunity, we performed transcriptomic analyses of leaf tissues of 5-week-old WT and *Oscds5* mutant lines. RNA samples were sequenced on the DNBSEQ-T7 platform by the paired-end RNA-sequencing method. Approximate 6-Gb data were generated and were further analyzed with an established bioinformatic pipeline (Wang et al., 2009, Zhao et al., 2022). Briefly, the clean RNA-sequencing reads were mapped to the Nipponbare reference genome using HISAT2, and the read numbers of each gene were counted using featureCounts. Differential gene expression analyses between samples were performed with the DESeq2 R package. Using the Benjamin-Hochberg method, we defined differentially expressed genes (DEGs) with the parameters as log2(fold change) > 1 and the false discovery rate (FDR) *p*-value < 0.05. Subsequently, 992 upregulated and 598 downregulated DEGs were identified in the *Oscds5* mutant compared to the WT (Figure 4a). The expression levels of selected DEGs were further validated with qRT-PCR assays (Figure 4c and Table S1). Among upregulated DEGs, many are involved in disease resistance, including *BSR1, CEBiP, Os2H16, OsNPR1, OsPBZ14, OsPR1b, OsWAK14*, and *PIBP1. BSR1* encodes a receptor-like cytoplasmic kinase and positively regulates innate immunity of rice. Overexpression of *BSR1* confers rice resistance to blast and bacterial blight (Sugano et al., 2018). *CEBiP* encodes a chitin-binding protein and acts as a positive regulator of chitin-triggered immunity (Akamatsu et al., 2013). The upregulated expression level of *CEBiP* is consistent with enhanced ROS production triggered by chitin in the *Oscds5* mutant. *Os2H16* encodes a short-chain peptide, a contributor to plant immunity and drought stress responses (Li et al., 2013). OsNPR1, a common positive regulator of plant immunity, plays an important role in the SA signaling pathway (Zhang et al., 2023). OsWAK14 is a wall-associated kinase and positively regulates rice blast resistance (Delteil et al., 2016). PIBP1, an RNA recognition motif (RRM)-containing protein, binds to the promoters of *OsWAK14* and *OsPAL1* to promote their expression levels and enhance rice blast resistance (Zhai et al., 2019). Gene Ontology (GO) enrichment analyses of DEGs were performed with TBtools (Chen et al., 2023). A total of 12 GO categories of upregulated DEGs were enriched in mutant *Oscds5*, including genes involved in responses to stress, biotic/abiotic stimuli and metabolic processes (Figure 4b).

**FIGURE 4.**
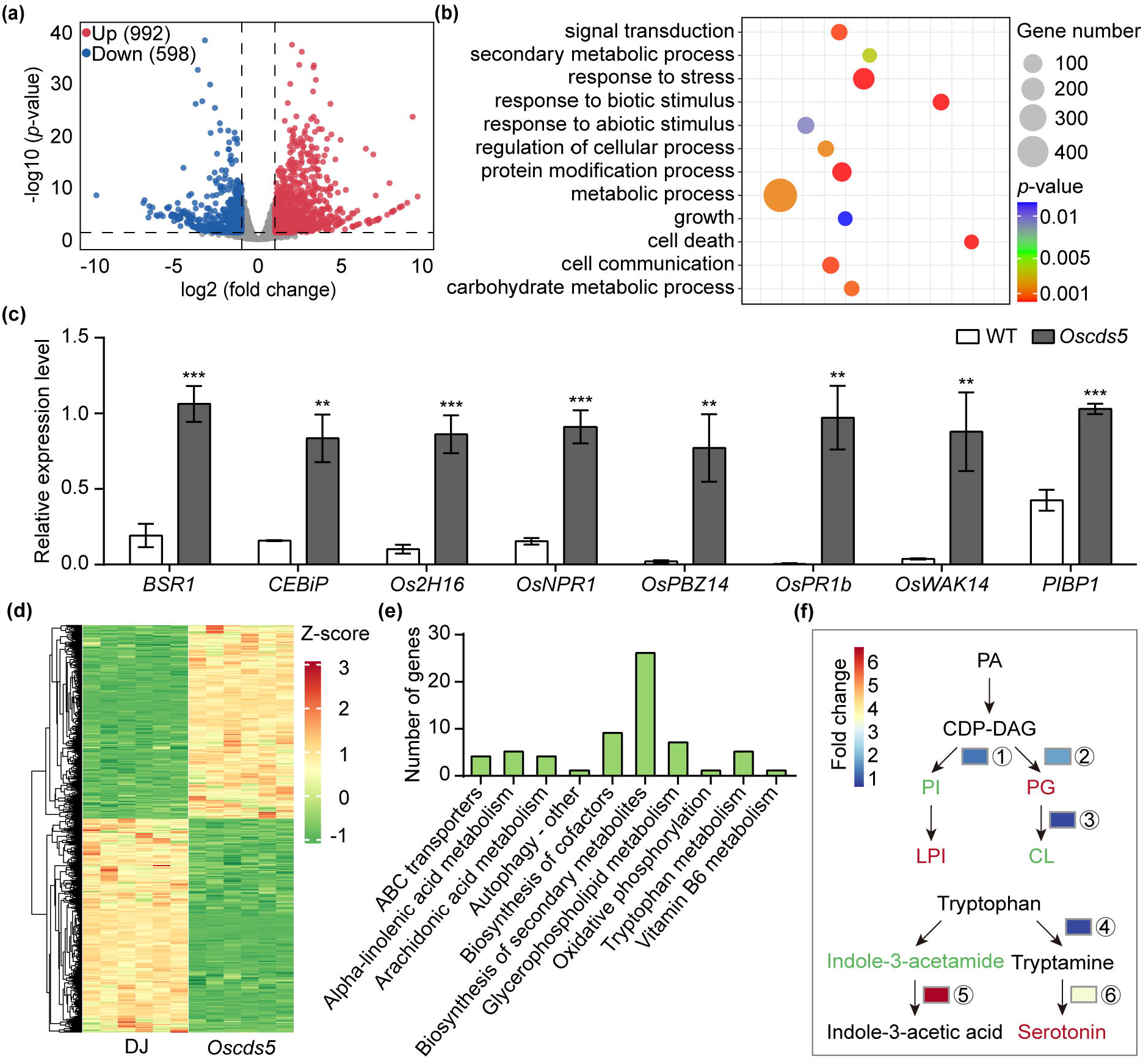
Integrated transcriptomic and metabolomic analyses of the *Oscds5* mutant. (a) Volcano plots of differentially expressed genes (DEGs) in transcriptomic analyses. The *x*-axis indicates the fold change in gene expression between the WT and mutant *Oscds5*, and the *y-*axis indicates statistically significant differences. Leaves of 5-week-old rice plants were used in the transcriptomic analyses. (b) Gene ontology (GO) analysis of upregulated DEGs in mutant *Oscds5*. (c) qRT-PCR assays of selected upregulated DEGs in mutant *Oscds5* (*n* = 3). (d) Heatmap of differential metabolites (*n* = 6). (e) KEGG enrichment analyses of differential metabolites. (f) A simplified metabolic flow scheme with the focus on phosphatidic acid (PA) metabolism and tryptophan metabolism. CDP-DAG, cytidinediphosphate-diacylglycerol; CL, cardiolipin; LPI, lyso-phosphatidylinositol; PG, phosphatidylglycerol; ①, PI synthase; ②, phosphatidylglycerolphosphate synthase; ③, cardiolipin synthase; ④, tryptophan decarboxylase; ⑤, indole-3-acetamide amidohydrolase; ⑥, tryptamine 5-hydroxylase. Red indicates metabolites with increased levels and green indicates metabolites decreased levels. *n* indicates the number of biological replicates used in the analyses. Asterisks indicate significant differences using the unpaired Student’s *t*-test (\*\**p* < 0.01, \*\*\**p* < 0.001).

To assess the metabolic changes in the *Oscds5* mutants, primary and secondary metabolites were analyzed with the non-target metabolomic technology on the liquid chromatography mass-spectroscopy (LC-MS) platform, and a total of 3,265 metabolites were detected. The differential metabolites were defined with parameters |Log2FC| ≥ 1.0 and *p*-value < 0.05 compared with the WT. A total of 333 upregulated and 369 downregulated metabolites were identified, which were also clustered among different biological replicates, indicating good homogeneity of the biological replicates and the reliability of the metabolic data (Figure 4d). The KEGG enrichment analyses showed that differential metabolites were enriched in secondary metabolite metabolism, cofactor metabolism, glycerophospholipid metabolism and tryptophan metabolism (Figure 4e). Lipid metabolism and tryptophan metabolism are associated with disease resistance (Gao et al., 2017, Wolinska et al., 2021). As shown in the transcriptome analyses, the genes in these pathways were mostly upregulated, such as for genes involved in PG and serotonin biosynthesis (Figure 4f). The elevated level of serotonin was conducive to enhanced disease resistance of rice (Jin et al., 2015). In addition, we found that several metabolites associated with biotic and abiotic stresses were also increased in the *Oscds5* mutant, including glyceollin III and raffinose. Glyceollin III boosts disease resistance to *Phytophthora sojae* in soybean (Jahan et al., 2020). Accumulation of raffinose enhances drought tolerance in maize and *A. thaliana* (Li et al., 2020). These transcriptomic and metabolomic results further show that mutation of *OsCDS5* enhances rice immunity and that OsCDS5 acts as a negative regulator of rice immunity.

The rice mutant *Oscds5* expresses broad-spectrum disease resistance and is altered in phospholipid metabolism. The PA levels increased, while PG and PI levels decreased in mutant *Oscds5*. Mutation of *OsCDS5* results in increased levels of ROS, defense-related genes and SA (Hong et al., 2018). All the phenotypes of mutant *Oscds5* are reminiscent of the *rbl1* mutant. In addition, *A. thaliana cds1 cds2* RNAi line shows enhanced resistance to oomycetes (Sha et al., 2023). Mutation of *RBL1* also results in reduced levels of PI and PI(4,5)P2, which are important for plant-fungal interactions, likely facilitating pathogen infection in rice. The level of PI is reduced in the *Oscds5* mutant, which may confer rice resistance to *M. oryzae*. As PI is a precursor of PI(4,5)P2, the level of PI(4,5)P2 in mutant *Oscds5* likely decreases, but needs to be investigated further. Mutant *Oscds5* has pale leaves at the seedling stage but no obvious growth defects later. In contrast, the *rbl1* line shows severe growth defects, spontaneous disease lesions and cell death, and the *Atcds1 Atcds2* mutant is lethal (Zhou et al., 2013). These results together indicate that OsCDS5 plays a relatively minor role in plant growth.

The function of plant CDP-DAG synthases including OsCDS5 and relevant genetic engineering of *CDSs* are worthy of further investigation. OsCDS5 is a negative regulator of biotic and abiotic stress responses, a promising candidate for crop engineering for resilience to biotic and abiotic stresses. Additionally, negative regulators are particularly compatible with genome editing technologies that are efficient in generating complete or partial loss-of-function alleles (Zhan et al., 2021), which could confer resilience of crops to biotic and abiotic stresses without compromising the yield. For example, genome editing of *RBL1* and disease-susceptibility genes *MLO* confers crops broad-spectrum disease resistance without yield losses (Li et al., 2022, Sha et al., 2023). Genome editing of *OsPQT3*, an E3 ubiquitin ligase-coding gene of rice, confers enhanced tolerance to salt stress and improved grain yield (Alfatih et al., 2020). Another strategy is to use the upstream open reading frames (uORFs), *cis*-regulatory elements in the promoter, which regulate protein translation (Xue et al., 2023). The *A. thaliana* snc1 is an autoactivated nucleotide binding-leucine rich repeat (NBS-LRR) immune receptor, and AtNPR1 is a positive regulator in plant immune (Xu et al., 2017). By inserting one uORF, the expression of which is specifically induced by pathogen infection, into the promoter of snc1 and AtNPR1 to fine-tune their protein translation levels, Xu et al. (2017) engineered *A. thaliana* and rice plants with broad-spectrum disease resistance without compromising the yield. In addition, using RNA interference (RNAi) technology to regulate gene expression levels, the *OsMADS26* RNAi rice plants show enhanced resistance to *M. oryzae, Xoo* and drought without yield losses. Using similar strategies with stress responsive regulatory elements, we may be able to engineer rice *OsCDS5* to achieve improved hyperosmotic tolerance and broad-spectrum disease resistance without significant growth trade-offs.

## ACKNOWLEDGEMENTS

We thank Prof. Y. Hong for providing rice seeds of DJ, mutant *Oscds5* and complementation lines; Prof. L. Dunkle for critical reading of the manuscript. Luminescence data were acquired at the National Key Laboratory of Agricultural Microbiology Core Facility. This work was supported by National Key R&D Program of China (2022YFA1304402), National Natural Science Foundation of China (32172373) and Fundamental Research Funds for the Central Universities (2023ZKPY002, 2662023PY006 and AML2023A05) to G. L. This work was also supported by Hubei Hongshan Laboratory.

## DATA AVAILABILITY STATEMENT

The gene ID for OsCDS5 is LOC_Os01g61560, and its accession number AP014957.1. RNA-seq data are available in the National Genomics Data Center (NGDC)’s Genome Sequence Archive (GSA) (Chen et al., 2021) and the National Center for Biotechnology Information (NCBI) (http://www.ncbi.nlm.nih.gov/) under the accession numbers PRJCA020630 and PRJNA1029920 and, respectively.

## REFERENCES

Akamatsu, A., Wong, H. L., Fujiwara, M., Okuda, J., Nishide, K., Uno, K., et al. (2013) An OsCEBiP/OsCERK1-OsRacGEF1-OsRac1 module is an essential early component of chitin-induced rice immunity. Cell Host & Microbe, 13, 465–476.

Alfatih, A., Wu, J., Jan, S. U., Zhang, Z. S., Xia, J. Q. and Xiang, C. B. (2020) Loss of rice PARAQUAT_TOLERANCE 3 confers enhanced resistance to abiotic stresses and increases grain yield in field. Plant, Cell & Environment 43, 2743–2754.

Blunsom, N. J. and Cockcroft, S. (2020) CDP-Diacylglycerol Synthases (CDS): gateway to phosphatidylinositol and cardiolipin synthesis. Frontiers in Cell and Developmental Biology, 8, 63.

Chen, C., Wu, Y., Li, J., Wang, X., Zeng, Z., Xu, J., et al. (2023) TBtools-II: a “one for all, all for one” bioinformatics platform for biological big-data mining. Molecular Plant, S1674-2052(1623)00281–00282.

Chen, T., Chen, X., Zhang, S., Zhu, J., Tang, B., Wang, A., et al. (2021) The Genome Sequence Archive Family: Toward Explosive Data Growth and Diverse Data Types. Genomics Proteomics Bioinformatics, 19, 578–583.

Choi, N., Im, J. H., Lee, E., Lee, J., Choi, C., Park, S. R., et al. (2020) WRKY10 transcriptional regulatory cascades in rice are involved in basal defense and Xa1-mediated resistance. Journal Of Experimental Botany, 71, 3735–3748.

Delteil, A., Gobbato, E., Cayrol, B., Estevan, J., Michel-Romiti, C., Dievart, A., et al. (2016) Several wall-associated kinases participate positively and negatively in basal defense against rice blast fungus. BMC Plant Biology, 16, 17.

Dhankher, O. P. and Foyer, C. H. (2018) Climate resilient crops for improving global food security and safety. Plant, Cell & Environment, 41, 877–884.

Gao, M., Yin, X., Yang, W., Lam, S. M., Tong, X., Liu, J., et al. (2017) GDSL lipases modulate immunity through lipid homeostasis in rice. PLoS Pathog, 13, e1006724.

Haselier, A., Akbari, H., Weth, A., Baumgartner, W. and Frentzen, M. (2010) Two closely related genes of Arabidopsis encode plastidial cytidinediphosphate diacylglycerol synthases essential for photoautotrophic growth. Plant Physiology, 153, 1372–1384.

Hong, Y., Yuan, S., Sun, L., Wang, X. and Hong, Y. (2018) Cytidinediphosphate-diacylglycerol synthase 5 is required for phospholipid homeostasis and is negatively involved in hyperosmotic stress tolerance. The Plant Journal 94, 1038–1050.

Jahan, M. A., Harris, B., Lowery, M., Infante, A. M., Percifield, R. J. and Kovinich, N. (2020) Glyceollin Transcription Factor GmMYB29A2 Regulates Soybean Resistance to Phytophthora sojae. Plant Physiol, 183, 530–546.

Jennings, W. and Epand, R. M. (2020) CDP-diacylglycerol, a critical intermediate in lipid metabolism. Chemistry and Physics of Lipids, 230, 104914.

Jin, B., Zhou, X., Jiang, B., Gu, Z., Zhang, P., Qian, Q., et al. (2015) Transcriptome profiling of the spl5 mutant reveals that SPL5 has a negative role in the biosynthesis of serotonin for rice disease resistance. Rice (N Y), 8, 18.

Kawano, Y., Akamatsu, A., Hayashi, K., Housen, Y., Okuda, J., Yao, A., et al. (2010) Activation of a Rac GTPase by the NLR family disease resistance protein Pit plays a critical role in rice innate immunity. Cell Host & Microbe, 7, 362–375.

Ke, Y., Hui, S. and Yuan, M. (2017) Xanthomonas oryzae pv. oryzae inoculation and growth rate on rice by leaf clipping method. Bio-protocol 7, e2568.

Li, N., Kong, L., Zhou, W., Zhang, X., Wei, S., Ding, X., et al. (2013) Overexpression of Os2H16 enhances resistance to phytopathogens and tolerance to drought stress in rice. Plant Cell, Tissue and Organ Culture, 115, 429–441.

Li, S., Lin, D., Zhang, Y., Deng, M., Chen, Y., Lv, B., et al. (2022) Genome-edited powdery mildew resistance in wheat without growth penalties. Nature, 602, 455–460.

Li, T., Zhang, Y., Liu, Y., Li, X., Hao, G., Han, Q., et al. (2020) Raffinose synthase enhances drought tolerance through raffinose synthesis or galactinol hydrolysis in maize and Arabidopsis plants. J Biol Chem, 295, 8064–8077.

Li, W., Chern, M., Yin, J., Wang, J. and Chen, X. (2019) Recent advances in broad-spectrum resistance to the rice blast disease. Current Opinion in Plant Biology, 50, 114–120.

Li, W., Zhu, Z., Chern, M., Yin, J., Yang, C., Ran, L., et al. (2017) A natural allele of a transcription factor in rice confers broad-spectrum blast resistance. Cell, 170, 114–126.e115.

Liu, J., Nie, B., Yu, B., Xu, F., Zhang, Q., Wang, Y., et al. (2023) Rice ubiquitin-conjugating enzyme OsUbc13 negatively regulates immunity against pathogens by enhancing the activity of OsSnRK1a. Plant Biotechnology Journal, 21, 1590–1610.

Liu, W., Liu, J., Triplett, L., Leach, J. E. and Wang, G. L. (2014) Novel insights into rice innate immunity against bacterial and fungal pathogens. Annual Review of Phytopathology, 52, 213–241.

Liu, Z., Zhu, Y., Shi, H., Qiu, J., Ding, X. and Kou, Y. (2021) Recent progress in rice broad-spectrum disease resistance. International Journal of Molecular Sciences, 22.

Ma, J., Morel, J. B., Riemann, M. and Nick, P. (2022) Jasmonic acid contributes to rice resistance against Magnaporthe oryzae. BMC Plant Biology, 22, 601.

Mei, C., Qi, M., Sheng, G. and Yang, Y. (2006) Inducible overexpression of a rice allene oxide synthase gene increases the endogenous jasmonic acid level, PR gene expression, and host resistance to fungal infection. Molecular Plant-Microbe Interactions, 19, 1127–1137.

Nalley, L., Tsiboe, F., Durand-Morat, A., Shew, A. and Thoma, G. (2016) Economic and environmental impact of rice blast pathogen (Magnaporthe oryzae) alleviation in the United States. PLoS One, 11, e0167295.

Qin, L., Zhou, Z., Li, Q., Zhai, C., Liu, L., Quilichini, T. D., et al. (2020) Specific recruitment of phosphoinositide species to the plant-pathogen interfacial membrane underlies Arabidopsis susceptibility to fungal infection. The Plant Cell, 32, 1665–1688.

Sha, G., Sun, P., Kong, X., Han, X., Sun, Q., Fouillen, L., et al. (2023) Genome editing of a rice CDP-DAG synthase confers multipathogen resistance. Nature, 618, 1017–1023.

Shen, H., Heacock, P. N., Clancey, C. J. and Dowhan, W. (1996) The CDS1 gene encoding CDP-diacylglycerol synthase in Saccharomyces cerevisiae is essential for cell growth. Journal of Biological Chemistry 271, 789–795.

Song, Y., Chong-Rui, A., Shao-Juan, J. and Di-Qiu, Y. (2010) Research progress on functional analysis of rice WRKY genes. Rice Science, 17, 60–72.

Sugano, S., Maeda, S., Hayashi, N., Kajiwara, H., Inoue, H., Jiang, C. J., et al. (2018) Tyrosine phosphorylation of a receptor-like cytoplasmic kinase, BSR1, plays a crucial role in resistance to multiple pathogens in rice. The Plant Journal, 96, 1137–1147.

Tang, B., Liu, C., Li, Z., Zhang, X., Zhou, S., Wang, G. L., et al. (2021) Multilayer regulatory landscape during pattern-triggered immunity in rice. Plant Biotechnology Journal 19, 2629–2645.

Tonnessen, B. W., Manosalva, P., Lang, J. M., Baraoidan, M., Bordeos, A., Mauleon, R., et al. (2015) Rice phenylalanine ammonia-lyase gene OsPAL4 is associated with broad spectrum disease resistance. Plant Molecular Biology, 87, 273–286.

Wang, Z., Gerstein, M. and Snyder, M. (2009) RNA-Seq: a revolutionary tool for transcriptomics. Nature Reviews Genetics, 10, 57–63.

Wolinska, K. W., Vannier, N., Thiergart, T., Pickel, B., Gremmen, S., Piasecka, A., et al. (2021) Tryptophan metabolism and bacterial commensals prevent fungal dysbiosis in Arabidopsis roots. Proc Natl Acad Sci U S A, 118.

Xu, G., Yuan, M., Ai, C., Liu, L., Zhuang, E., Karapetyan, S., et al. (2017) uORF-mediated translation allows engineered plant disease resistance without fitness costs. Nature, 545, 491–494.

Xue, C., Qiu, F., Wang, Y., Li, B., Zhao, K. T., Chen, K., et al. (2023) Tuning plant phenotypes by precise, graded downregulation of gene expression. Nature Biotechnology.

Yang, L., Zhao, M., Sha, G., Sun, Q., Gong, Q., Yang, Q., et al. (2022) The genome of the rice variety LTH provides insight into its universal susceptibility mechanism to worldwide rice blast fungal strains. Computational and Structural Biotechnology Journal, 20, 1012–1026.

Yang, W., Liu, X., Liu, M., Wei, F., Yang, L., Yuan, M., et al. (2023) High-quality complete genome sequence of Xanthomonas oryzae pv. oryzicola (Xoc) strain HB8. Microbiology Resource Announcements, 12, e0045923.

Yang, Y., Lee, M. and Fairn, G. D. (2018) Phospholipid subcellular localization and dynamics. Journal of Biological Chemistry, 293, 6230–6240.

Zhai, K., Deng, Y., Liang, D., Tang, J., Liu, J., Yan, B., et al. (2019) RRM transcription factors interact with NLRs and regulate broad-spectrum blast resistance in rice. Molecular Cell, 74, 996–1009.e1007.

Zhan, X., Lu, Y., Zhu, J.-K. and Botella, J. R. (2021) Genome editing for plant research and crop improvement. Journal of Integrative Plant Biology, 63, 3–33.

Zhang, H., Wang, F., Song, W., Yang, Z., Li, L., Ma, Q., et al. (2023) Different viral effectors suppress hormone-mediated antiviral immunity of rice coordinated by OsNPR1. Nature Communications, 14, 3011.

Zhao, J., Sun, P., Sun, Q., Li, R., Qin, Z., Sha, G., et al. (2022) The MoPah1 phosphatidate phosphatase is involved in lipid metabolism, development, and pathogenesis in Magnaporthe oryzae. Molecular Plant Pathology, 23, 720–732.

Zhou, Y., Peisker, H., Weth, A., Baumgartner, W., Dörmann, P. and Frentzen, M. (2013) Extraplastidial cytidinediphosphate diacylglycerol synthase activity is required for vegetative development in Arabidopsis thaliana. The Plant Journal, 75, 867–879.

