## Supplemental Table 1 for "Mutation of *OsCDS5* confers broad-spectrum disease resistance in rice"

**Supporting information**

**Table S1 Primers used in this study**

| Name | Forward Primer (5’-3’) | Reverse Primer (5’-3’) |
| --- | --- | --- |
| qRT-Actin/FR | CAGGCCGTCCTCTCTCTGTA | AAGGATAGCATGGGGGAGAG |
| qRT-BSR1/FR | CCGGGACTTCAAAGCATCTAAC | TGTTGGTCCCTCCCTTGCT |
| qRT-CEBiP/FR | CGTGTTCAACGCCTTCGT | CGTCTGGCTGACATTTATCTTG |
| qRT-Os2H16/FR | CAGTGGCGAGGCGAGTAAG | GTGTTTGGGCTTGGTGTTTATG |
| qRT-OsAOS2/FR | AAGCTGCTGCAATACGTGTACTGG | CGACGAGCAACAGCCTTCCG |
| qRT-OsNPR1/FR | ATCTTATCTGTTGCCAACTTATGC | CAAGGTTTGACCGGACTACCA |
| qRT-OsPAL1/FR | CTACCCGCTGATGAAGAAGC | GAACCTTGTTCAGCTCCTCG |
| qRT-OsPBZ14/FR | ATGAAGCTCAACCCTGCTGT | TGAGCTTGCCCACCTTACTT |
| qRT-OsPR1a/FR | CGTGTCGGCGTGGGTGT | GGCGAGTAGTTGCAGGTGATG |
| qRT-OsPR1b/FR | TACGCCAGCCAGAGGAGC | GCCGAACCCCAGAAGAGG |
| qRT-OsWAK14/FR | CCGTTCTGAACAGGGTATGC | CTTTGCCACCTGGCCTTAAT |
| qRT-OsWRKY70/FR | CCGCTGCTGTTTTGATCATCT | GGAGCTAAGCTAACTCACTCCACA |
| qRT-PIBP/FR | TGTGAACTATCAACTGCCTCC | CATCCTCTGCCTTCCTGACTA |
| qRT-Pot2/FR | ACGACCCGTCTTTACTTATTTGG | AAGTAGCGTTGGTTTTGTTGGAT |
| qRT-Ubi/FR | TTCTGGTCCTTCCACTTTCAG | ACGATTGATTTAACCAGTCCATGA |
| qRT-Xoc/FR | CAGTTGCGTGCGTAGTTCAG | ACAGCAGTTCGATGCCTACA |
| qRT-Xoo/FR | CCAGCCGTTGCGTTTGTC | TCAGCATGGTCGCCGTAGAA |
